# Whole Genome Variation of Transposable Element Insertions in a Maize Diversity Panel

**DOI:** 10.1101/2020.09.25.314401

**Authors:** Yinjie Qiu, Christine H. O’Connor, Rafael Della Coletta, Jonathan S. Renk, Patrick J. Monnahan, Jaclyn M. Noshay, Zhikai Liang, Amanda Gilbert, Sarah N. Anderson, Suzanne E. McGaugh, Nathan M. Springer, Candice N. Hirsch

## Abstract

Intact transposable elements (TEs) account for 65% of the maize genome and can impact gene function and regulation. Although TEs comprise the majority of the maize genome and affect important phenotypes, genome wide patterns of TE polymorphisms in maize have only been studied in a handful of maize genotypes, due to the challenging nature of assessing highly repetitive sequences. We implemented a method to use short read sequencing data from 509 diverse inbred lines to classify the presence/absence of 445,418 non-redundant TEs that were previously annotated in four genome assemblies including B73, Mo17, PH207, and W22. Different orders of TEs (i.e. LTRs, Helitrons, TIRs) had different frequency distributions within the population. LTRs with lower LTR similarity were generally more frequent in the population than LTRs with higher LTR similarity, though high frequency insertions with very high LTR similarity were observed. LTR similarity and frequency estimates of nested elements and the outer elements in which they insert revealed that most nesting events occurred very near the timing of the outer element insertion. TEs within genes were at higher frequency than those that were outside of genes and this is particularly true for those not inserted into introns. Many TE insertional polymorphisms observed in this population were tagged by SNP markers. However, there were also 19.9% of the TE polymorphisms that were not well tagged by SNPs (R^2^ < 0.5) that potentially represent information that has not been well captured in previous SNP based marker-trait association studies. This study provides a population scale genome-wide assessment of TE variation in maize, and provides valuable insight on variation in TEs in maize and factors that contribute to this variation.

## INTRODUCTION

Transposable elements (TEs) are present in all eukaryotic genomes (Bennetzen 2000; Wicker *et al.* 2007). In maize, 65% of the genome is made up of intact TEs (Jiao *et al.* 2017), and another 20% is comprised of fragmented TEs (Schnable *et al.* 2009). There are many examples of phenotypic effects of TEs from null mutations, such as maize kernel color (Selinger and Chandler 2001), white wine grapes (Cadle-Davidson and Owens 2008), and color variation in the common morning glory (Clegg and Durbin 2000). TE insertions can also positively or negatively affect gene regulatory functions, such as insertion of a long terminal repeat LTR retrotransposon into the promoter region of the Ruby gene in oranges that leads to red fruit flesh of blood oranges (Butelli *et al.* 2012), and an LTR retrotransposon that is associated with red skin color in apples (Zhang *et al.* 2019). TE insertions in maize have also been associated with genes that are upregulated in response to abiotic stress (Makarevitch *et al.* 2015).

TEs are classified into two classes depending on how they replicate and from there into superfamilies and families by sequence similarity (Wicker *et al.* 2007). Class I elements, or retrotransposons, replicate via an RNA intermediate (Bennetzen 2000; Lisch 2013). Long terminal repeat retrotransposons (LTR) are the most abundant type of retrotransposons in maize (Bennetzen 2000) and intact elements account for over half of the maize genome by sequence length (Anderson *et al.* 2019; Stitzer *et al.* 2019). Class II elements, or DNA transposable elements, replicate via a DNA intermediate, and the two largest orders are terminal inverted repeat (TIR) and Helitron elements. TIRs are defined by terminal inverted repeat sequences at both ends of the TE (Wicker *et al.* 2007) and intact TIRs make up around 3% of the maize genome (Anderson *et al.* 2019). Helitrons are defined by their ‘rolling circle’ replication mechanism (Lisch 2013) and intact Helitrons make up around 4% of the maize genome (Anderson *et al.* 2019). TEs are found throughout the maize genome, they can be found near and even within genes, can be anywhere from a few hundred base pairs to >10 kb in length (Bennetzen 2000), and range in age from very recent insertions to >2 million years old insertions (Stitzer *et al.* 2019).

Studies done on TEs in maize have shown extensive variation in TE insertion presence/absence patterns at specific loci across maize inbred lines (Sanmiguel *et al.* 1996; Fu and Dooner 2002; Morgante *et al.* 2005; Dooner *et al.* 2019). Early work on the *bronze* locus in multiple maize lines found that different lines differed in not only the gene order and content but also in TE content (Fu and Dooner 2002). More recent work on mutations in the same region found that not only were high mutation rates due to TE insertions, but that different TEs were inserting in different maize lines (Dooner *et al.* 2019). Whole genome analysis of four maize genomes with *de novo* TE annotations revealed extensive TE polymorphism between maize lines on a whole genome scale (Anderson *et al.* 2019). On average, about 500 Mb of TE sequence, or ~20% of the maize genome, was variable between the four inbred lines (B73, Mo17, PH207 and W22). Another 1.6 Gb of TE sequence was only shared between two or three of the lines.

Genome-wide TE presence/absence polymorphism at a population scale in crop species has recently been investigated using short reads whole genome resequencing data in a number of species. For example, by sequencing 602 cultivated and wild tomato accessions, Domínguez et al identified at least 40 TE polymorphism that were not tagged by SNPs, and were associated with traits such as fruit color (Dominguez *et al.* 2020). Another example is the resequencing of 3,000 Asian rice varieties, which identified polymorphic TEs at low frequency that were associated with rice domestication (Carpentier *et al.* 2019). Despite the widespread prevalence and polymorphism of TEs in the maize pan-genome, as well as many examples connecting TE insertions to functional phenotypic variation, there have been very few scans of specific TE insertion frequencies in divergent maize populations. The analysis of the frequency of TE insertions can provide insights into the level of variability for TEs and help understand the presence/absence of common and rare TE variants. To understand patterns of TE polymorphism on a genome-wide scale, we utilized short read sequencing of 509 diverse maize lines to score the presence/absence of 445,418 non-redundant TEs that were annotated in four reference genome assemblies (B73, Mo17, PH207, and W22). This study provides a genome-wide analysis of TE presence/absence polymorphism across a large panel of diverse maize genotypes as we continue to try to understand how TEs contribute to phenotypic variation and adaptation within the species.

## MATERIALS AND METHODS

### Whole genome resequencing

A subset of 509 lines from the Wisconsin Diversity Panel (Hansey *et al.* 2010; Hirsch *et al.* 2014; Mazaheri *et al.* 2019) was used for this study (Table S1). For 57 genotypes, available short-read sequence data was downloaded from the National Center for Biotechnology Information (NCBI) Sequence Read Archive (SRA; Table S1). These samples ranged in theoretical coverage of 10-55x sequencing depth based an estimated genome size of 2.4 Gb. For 454 genotypes, plants were grown under greenhouse conditions (27C/24C day/night and 16 h light/8 h dark) with five plants of a single genotype per pot. Plants were grown in Metro-Mix 300 (Sun Gro Horticulture) with no additional fertilizer. Tissue was harvested for DNA extractions at the Vegetative 2 developmental stage. The newest leaf of each seedling in the pot was collected and immediately flash frozen in liquid nitrogen. Tissue was ground in liquid nitrogen using a mortar and pestle. DNA was extracted using a standard cetyltrimethylammonium bromide (CTAB) DNA extraction protocol (Saghai-Maroof *et al.* 1984), and treated with 25 uL of PureLink RNase A (Invitrogen) at 39°C for 30 minutes. Genomic DNA for each genotype was submitted to Novogene (Novogene Co., Ltd., Beijing, China) for whole genome sequencing with 150 base pair paired end reads generated on a HiSeq X Ten sequencing machine. For each genotype at least 20x theoretical sequencing depth was achieved.

### Read Alignment and processing

Quality control analysis of the sequence data was conducted using fastqc version 0.11.7 (https://www.bioinformatics.babraham.ac.uk/projects/fastqc/). Adapter sequence and low-quality base trimming was done using cutadapt version 1.18 (Martin 2011) and sickle version 1.33 (https://github.com/najoshi/sickle) both with default parameters. Sequence reads were aligned to the B73 v4 (Jiao *et al.* 2017), Mo17 v1 (Sun *et al.* 2018), PH207 v1 (Hirsch *et al.* 2016), and W22 v1 (Springer *et al.* 2018) genome assemblies. Alignment of reads was conducted using SpeedSeq version 0.1.2 (Chiang *et al.* 2015), which efficiently parallelizes bwa mem (Li and Durbin 2009), with read groups labelled separately for each FASTQ. Alignments were subsequently filtered to require a minimum mapping quality of 20 using samtools view (Li *et al.* 2009). To assess sample integrity, single nucleotide polymorphisms (SNPs) were called relative to the B73 v4 genome assembly using Platypus v 0.8.1 (Rimmer *et al.* 2014) with default parameters. A subset of 50,000 random bi-allelic SNPs with less than 50% missing data and less than 10% heterozygosity was used to calculate Nei’s pairwise genetic distances using StAMPP version 1.5.1 and a neighbor joining tree was generated with nj() in ape version 5.3 all using R version 3.6.2 (R Development Core Team 2020). Three samples with discordance to known pedigree relationships were removed from downstream analysis to result in the 509 genotypes included in Table S1.

### Transposable element identification in the Wisconsin Diversity Panel

Transposable element (TE) annotations for the B73, Mo17, PH207, and W22 genome assemblies were obtained from https://mcstitzer.github.io/maize_TEs. Mean coverage over the 10 bp window internal to the start and end of each TE was calculated using multicov within bedtools version 2.29.2 (Quinlan and Hall 2010) (Figure S1). Binary alignment map (BAM) files for B73, Mo17, PH207, and W22 short reads aligned to each genome assembly were down sampled to 15x and 30x coverage using sambamba version 0.8.0 (Tarasov *et al.* 2015) for model training. If coverage was at or below either of these thresholds no down sampling was performed (Table S2). The caret package in R version 3.6.3 (R Development Core Team 2020) was used to train a random forest model using the rf function and validated with 10 fold cross validation repeated three times. The training set consisted of counts from resequencing data from three of the four genomes aligned to all four reference genome assemblies and the test set consisted of the counts from resequencing data from the fourth genome mapped to all for reference genome assemblies. The mean coverage over the 10 bp at the start of the TE, mean coverage over the 10 bp at the end of the TE, and the TE order were used as predictors to train the model. Separate models were trained for realized coverage of 30x and realized coverage of 15x using the down sampled BAM files described above. Each model was trained with 500,000 features that were selected from the full set of features due to computational limitations. These features were selected to have approximately 50% present features and 50% absent features so as not to bias the model accuracy for presence or absence. A non-redundant set of TEs with presence/absence scores based on whole-genome comparisons of B73, Mo17, PH207, and W22 was used as previously reported (Anderson *et al.* 2019) for analysis of the true positive, true absent, false positive, and false absent rates of the short read based presence/absence calls.

A final model was trained for 15x coverage and 30x coverage using resequencing data from all four genomes mapped to all four genome assemblies as described above using 500,000 training features with either 15x or 30x coverage. These models were used to estimate the probability of presence of a TE based on the two internal coverage windows and the TE order for all of the resequenced genomes. If the realized coverage for a sample was >=25x depth the 30x model was used and if the realized coverage for a sample was <25x depth the 15x model was used (Figure S2). If the probability of present from the model was >=0.7 a TE was classified as present in the sample. If the probability of present from the model was <=0.3 a TE was classified as absent in the sample. All other TEs were classified as ambiguous. For any TE where the resequencing reads mapped to its cognate genome (e.g. B73 reads mapped to the B73 genome assembly) did not result in a present classification the TE was considered recalcitrant to accurate calls for short read data and was removed from downstream analysis. Across all samples mapped to a reference genome assembly if there was greater than 25% ambiguous data for a TE the TE was removed from downstream. Presence, absence, and ambiguous scores for each TE that was retained can be found in File S1 for B73, File S2 for Mo17, File S3 for PH207, and File S4 for W22.

The non-redundant TE dataset from Anderson et al 2019 was used to combine TE population frequencies across homologous TEs from the different genome assemblies. The mean of the population frequency for each TE was calculated between homologous TEs (File S5). TE metadata (e.g. location relative to the nearest gene or family) for the TE in the reference genome it was first identified in was used in downstream analyses using a previously described order for adding in TEs from each of the reference genome assemblies (Anderson *et al.* 2019).

Information about TE family/superfamily size and LTR similarity of LTR retrotransposons was obtained from the previously published TE annotation gff files (Anderson *et al.* 2019). Gene annotations from B73 version 4 (Jiao *et al.* 2017), Mo17 (Sun *et al.* 2018), PH207 (Hirsch *et al.* 2016), and W22 (Springer *et al.* 2018) reference genomes were used to identify TE locations relative to genes. All metadata is included in File S5.

The Kolmogorov-Smirnov test was used to test whether the frequency distributions of nested and non-nested TEs shown in Figure 1 were different. This test was implemented using the ks.test function in base R version 3.6.3 (R Development Core Team 2020).

**Figure 1.**
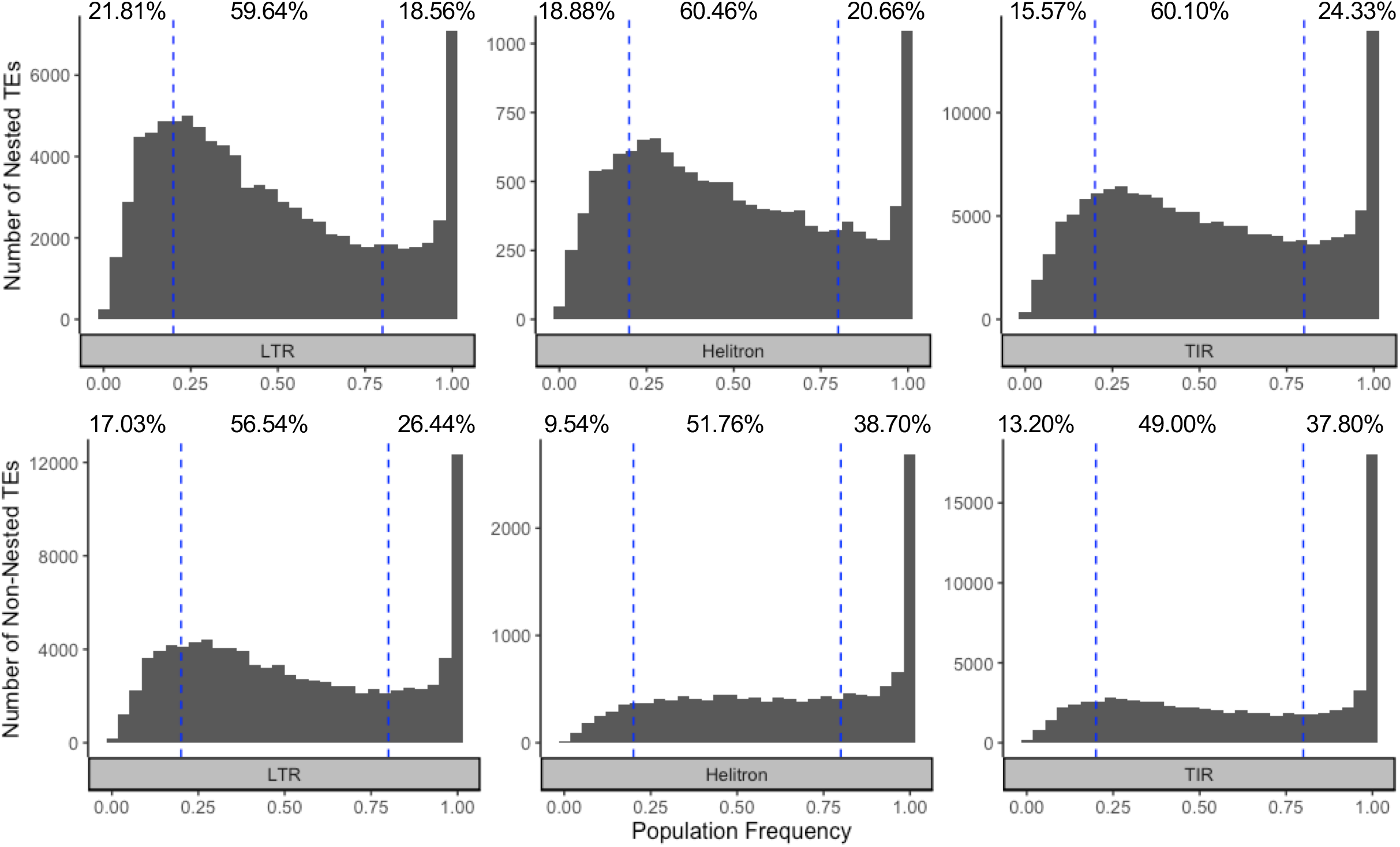
TE frequency distribution of a non-redundant set of TEs annotated in the B73, Mo17, PH207, and W22 genome assemblies. Short read sequence data from 509 genotypes were aligned to each genome assembly. Using a random forest machine learning method, TEs were classified into present (probability present >= 0.7), absent (probably present <=0.3), and all the other TEs were classified as ambiguous. For homologous TEs that were present in more than one assembly, the mean frequency across assemblies was calculated for the non-redundant set. TEs with less than 25% ambiguous calls are included (455,418 TEs). Percentages indicate the percent of low (<20%), moderate (20-80%), and high (>80%) frequency TEs in each order.

### SNP identification and analysis

Joint SNP calling was performed across all genotypes using *freebayes* v1.3.1-17 (Garrison and Marth 2012) for the alignments to the B73 v4 reference genome assembly with scaffolds removed. Sites with less than 1x average coverage over the whole population or greater than 2x the population level mean coverage were excluded from SNP calling. SNP calls were quality filtered using GATK (v4.1.2) (Van der Auwera *et al.* 2013) recommended filters as follows: QualByDepth (QD) less than 2, FisherStrand (FS) greater than 60, root mean square mapping quality (MQ) less than 40, MappingQualityRankSum (MQRankSum) less than −12.5, or ReadPosRankSum less than −8. If a SNP failed (e.g. MQ < 40) any one of those filters it was removed from the dataset. Finally, *vcftools* (v 0.1.13) (Danecek *et al.* 2011) was used to filter sites for a minimum quality score of 30, a minimum allele count of 50 (called in at least 25 homozygotes or 50 heterozygotes) and to filter out genotypes called in fewer than 90% of the individuals in the population. In total, 3,146,253 SNPs remained after all of these filtering steps (File S6).

These SNP calls were used to conduct linkage disequilibrium analysis with the TE presence/absence scores described above. For this analysis, TE presence/absence data from only the alignments to B73 reference genome assembly were used. Any SNPs located within TEs were removed. Linkage disequilibrium between SNPs and TEs was calculated using plink v1.90b6.16 (Purcell *et al.* 2007) with the --make-founders option to calculate LD among all inbred lines, --allow-extra-chr to calculate LD in extra scaffolds, --ld-window-r2 0 to report r^2^ in the 0 to 1 range (default is 0.2 to 1), --ld-window 1000000 --ld-window-kb 1000 to calculate LD within 1mb windows, and --r2 dprime with-freqs to report both D’ and r^2^ and to display minor allele frequencies in the output. If more than one SNP had the same highest LD in either the r^2^ or D’ analysis, the SNP that was physically closest to the SNP was used for downstream analyses. Principal components analysis of SNPs and TEs were conducted using Plink v1.90b6.18 (Purcell *et al.* 2007) with the -pca option. Pairwise genetic distance matrices for SNPs and TEs were calculated as 1 minus identity by state (IBS) using TASSEL version 5.2.64 (Bradbury *et al.* 2007).

### Data Availability

All sequence data is available on the NCBI SRA (BioProject PRJNA661271, Table S1). Code for this study is available at https://github.com/HirschLabUMN/TE_variation.

## RESULTS AND DISCUSSION

### Using short read sequence data to identify TE presence/absence

For this study, we implemented an approach for scoring TE presence/absence variation from short read sequence alignment using the average coverage of windows within the boundaries of previously annotated TEs in a random forest machine learning model. To assess classification accuracy, presence/absence scores defined by previous comparison of TE content generated for four maize genome assembles including B73, Mo17, PH207, and W22 were used as true positive (Anderson *et al.* 2019). It should be noted that some polymorphisms in this true positive set might be wrong for reasons described in the publication from which they were generated (Anderson *et al.* 2019). As such, 100% accuracy in comparison to this set is not possible unless the same miscalls are generated in two independent methods. Still, this set represents a high quality set of TE polymorphisms for which to assess the relative accuracy of different parameters. The average model accuracy observed across the training iterations was 0.88 (SE=0.01) for the model with 15x coverage and 0.89 (SE=0.04) for the model with 30x coverage. The final model used for prediction had prediction accuracies of 0.88 and 0.89 for 15x, 30x coverage model, respectively. The threshold for classification of present, absent, or ambiguous was selected to balance accuracy of the model across the different training sets and the proportion of TEs in which a non-ambiguous categorization could be assigned (Figure S3). For both the 15x and 30x models, a cutoff of <=0.3 probability of present to classify a TE as absent and a cutoff of >=0.7 probability of present to classify a TE as present was determined to optimally balance these two metrics.

These models and the subsequent filtering methods (i.e. removing TEs that are recalcitrant to short read based genotyping (B73 ref: 12.02%, Mo17: 5.60%, PH207: 1.32%, W22: 5.37%) and those with high levels of ambiguous calls (B73 ref: 0.20%, Mo17: 0.24%, PH207: 1.54%, W22: 0.34%)) provide a high quality set of TE presence/absence calls that allowed for analysis of TE variation on a genome-wide scale in maize. It should be noted, however, that rare alleles will be under-represented in this set due to the fact that only those TEs that were previously annotated in at least one of four *de novo* assembled maize genomes are included in this study.

### TE frequency distribution in a panel of diverse inbred lines

The final models were applied to short read sequence data from 509 genotypes of the Wisconsin Diversity Panel (Hansey *et al.* 2010; Hirsch *et al.* 2014; Mazaheri *et al.* 2019) mapped to the B73, Mo17, PH207, and W22 reference genome assemblies. For each annotated TE in these genome assemblies, the presence/absence frequency of the TE in this panel of diverse inbred lines was determined. Genetic relationships based on SNPs and TEs were assessed to further validate the quality of these TE presence/absence calls based on their consistency with other marker types and known pedigree relationships. Principle component analysis of the SNPs and TEs both revealed expected population structure based on previous pedigree information and heterotic group membership (Figure S4A), and pairwise genetic distances between individuals using SNPs and TEs were highly correlated (Figure S4B). As previously reported, there are a number of shared TEs across these four genome assemblies (Anderson *et al.* 2019). The population frequency of homologous TEs determined from mapping the short read data to the different genome assemblies were highly correlated (average Pearson’s correlation across the six pairwise comparisons r^2^ = 0.91 (SE=0.04); Figure S5), demonstrating the consistency of this pipeline, and enabling classifications to be combined into a non-redundant set of TEs. For downstream analysis of these redundant TEs, the mean was calculated across frequencies obtained from all genomes for which a homologous TE was present (Figure 1; File S5).

Different orders of TEs have different mechanisms of replication (Wicker *et al.* 2007), and families within these orders have different insertional preferences (SanMiguel *et al.* 1998; Sultana *et al.* 2017; Springer *et al.* 2018) and different effects on DNA methylation and chromatin accessibility (Eichten *et al.* 2012; Choi and Purugganan 2018; Noshay *et al.* 2019). As such, we hypothesized the frequency of TEs in the population would be variable across orders of TEs and families. For non-nested elements (i.e. those not contained within another TE), a subset of the TEs were present in all or nearly all of the diverse lines included in this study for LTRs, Helitrons, and TIRs (Figure 1). However, for all three orders, there was a substantial number of TEs that were at low (<20%) to moderate (20-80%) frequency in the population. This proportion was particularly high for the LTRs where 73.57% of non-nested TEs were present in low to moderate frequency in the population. These polymorphic TEs have the potential to drive phenotypic variation as has been seen for a number of morphological/developmental (Chuck *et al.* 2007; Studer *et al.* 2011) and adaptive traits (Yang *et al.* 2013) in maize. The demonstrated extent of TE variability on a genome-wide scale across a large number of individuals that this study provides is critical in furthering our understanding of the contribution of variable TEs in producing phenotypic variation within species. Given that only TEs that were annotated in at least one of only four genomes were included in this study, it is expected that the proportion of polymorphic TEs at low to moderate frequency would be substantially higher than what is presented here if all TEs in all genotypes in the population were annotated. In contrast, the majority of the high frequency TEs were likely already captured from just these four genome assemblies, and the count of these TEs would remain relatively static if all genomes were to be de novo assembled and annotated.

### LTR retrotransposons with lower similarity of their two LTRs were generally more frequent in the population than those with higher LTR similarity

A negative relationship between age of TE insertions approximated by the similarity of their two LTRs and their frequency was previously demonstrated using a limited number (n=4) of genotypes (Anderson *et al.* 2019). We sought to test this relationship between similarity of the two LTRs for an insertion and frequency on our broader set of germplasm, where younger insertions generally have higher LTR similarity and older insertions generally have lower LTR similarity. For other orders of TEs, there are not accurate methods to assess age. Thus, these analyses were limited to LTR retrotransposon insertions. In this maize diversity panel, the LTR similarity was negatively correlated with population frequency, which suggests that LTRs with low LTR similarity were generally more frequent in the population than LTRs with high LTR similarity (Figure 2a). It is worth noting, however, that there were a large number (n=14,509, 36.30% of all high frequency LTR) of LTR insertions that had moderately LTR similarity (LTR similarity 95-99%; n=12,655) or high LTR similarity (LTR similarity >99%; n=1,854), and are at high frequency (>80%) in the population (Figure 2b), and 180 of the insertions with high LTR similarity were fixed in this population. Fifteen of the fixed insertions with high LTR similarity were within 5kb of a gene and may be under selection or linked to other features in the genome under selection that could have driven their rapid rise to fixation in the population. A number of these genes have been functionally characterized, such as agpll2 (Huang *et al.* 2014), as well as trps4 (Zhou *et al.* 2014), which may be important for tolerance to different stress conditions.

**Figure 2.**
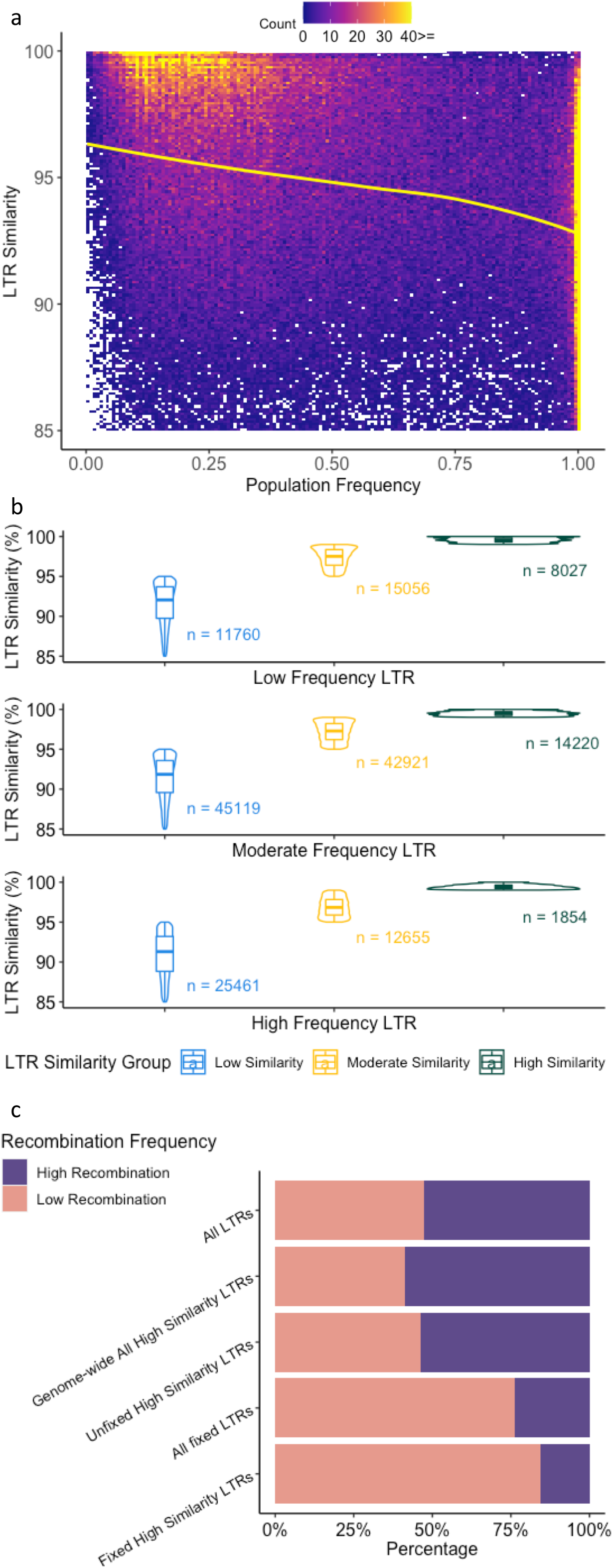
Relationship between TE similarity and frequency in a population of diverse inbred lines. a) Heatmap of LTR similarity versus frequency where white boxes indicate no TEs present at a particular frequency-by-LTR similarity. Yellow line is a LOESS curve fit through the data (n=177,073). b) Relationship of LTR similarity in categories of low similarity (LTR similarity <95%), moderate similarity (LTR similarity between 95-99%), and high similarity (LTR similarity >99%), and frequency in categories of low frequency (<20%), moderate frequency (20-80%), and high frequency (>80%). c) Proportion of different groups of LTRs in the low and high recombination portions of the genome based on B73 reference (n=108,968).

There was a large portion (84.52%) of the fixed insertions with high LTR similarity that were located within previously defined low recombination regions (Table S3; (Swanson-Wagner *et al.* 2010; Eichten *et al.* 2011)). These regions were defined by comparing genetic and physical maps and defining boundaries to define the high recombination arms and the low recombination middle of each chromosomes including the centromere and pericentromere. The fixed insertions with high LTR similarity were enriched (Fisher test with p-value < 0.001; Figure 2c) for being located within the low recombination regions of the genome relative to the frequency for all LTRs. However, this enrichment in the low recombination region of the genome is also observed for all fixed LTRs regardless of their LTR similarity. If we look at other elements in the families from which these fixed insertions with high LTR similarity are in there is no significant difference between them and all LTRs or all of the LTR insertions with high LTR similarity in the genome. Thus, the enrichment of fixed LTRs with high LTR similarity in the pericentromere is likely a product of their location.

The elements in the high frequency group (not just fixed) with high LTR similarity were from 494 different families. We sought to test if there was an enrichment within these families for elements that were at high frequency with high LTR similarity relative to the frequency of this class compared to all other LTRs. Indeed, for those families with at least 20 elements in the family (n=62), 15 had a higher than expected proportion of elements that had high LTR similarity and were at high frequency in the population (Fisher test with p-value < 0.01; Table S4). The majority of these enriched families are within the Gypsy superfamily, including RLG00001, a large Cinful-Zeon family (Sanz-Alferez *et al.* 2003) with 23,948 copies in the B73 genome that lacks homologs in Sorghum (Paterson *et al.* 2009; Jiao *et al.* 2017; Stitzer *et al.* 2019). Another family of note is RLG0009. This family was previously documented to be consistently upregulated under heat stress across genotypes with multiple members of the family showing increased expression, potentially due to the presence of conserved cis-regulatory elements within the TE that facilitate stress-responsive expression of this family (Liang *et al.* 2020). The potential importance of this family to stress responsiveness, the conserved response across members of the family, and the consistency in response across genotypes are all consistent with, and provide potential explanations for why, this family had enriched presence in the class of high frequency TEs that also have high LTR similarity.

On the other end of the spectrum, for the class of TEs with low LTR similarity (LTR similarity <95%), it is expected that some will be common as they have had time to rise in frequency in the population and other will be rare as they are being lost over time, which was the case in this population. Within the class of TEs that have low LTR similarity, 14.28% were at low frequency in the population, 30.92% were at high frequency in the population, and the remaining 54.80% were at moderate frequency in the population (Figure 2b).

### Most nested TEs insertions occur near the insertion time of the outer element

Nested TEs exhibited higher levels of moderate frequency (20-80%) and low frequency (<20%) TEs as compared to the non-nested TEs (Figure 1). In all three orders, the frequency distribution of nested and non-nested TEs was significantly different (KS test, p-value < 2×10-16). A nested TE could be the product of an insertion very soon after the outer element inserted or it could be the product of a spectrum of much younger insertions that happened well after the insertion of the outer element. Genome-wide we see that nested elements have higher LTR similarity than non-nested elements (Figure S6).

To further explore the difference in frequency of nested and non-nested elements we looked at specific nested TEs and the non-nested element into which they inserted. Within these pairs, the nested insertion should be at the same or lower frequency compared to the outer element, as the nested insertion cannot exist in a genotype without the outer element. This was true for 67.38% of the pairs, and an additional 20.33% had less than 5% higher frequency in the nested element. Overall, there was a strong correlation between the nested TE frequency and the frequency of the TE into which it was nested (Pearson’s correlation r^2^ = 0.52; Figure 3a), with 71.5% having frequencies within 5% of each other, and the remaining 28.5% of pairs having a range of difference in frequencies between the inner and outer elements (Figure 3b).

**Figure 3.**
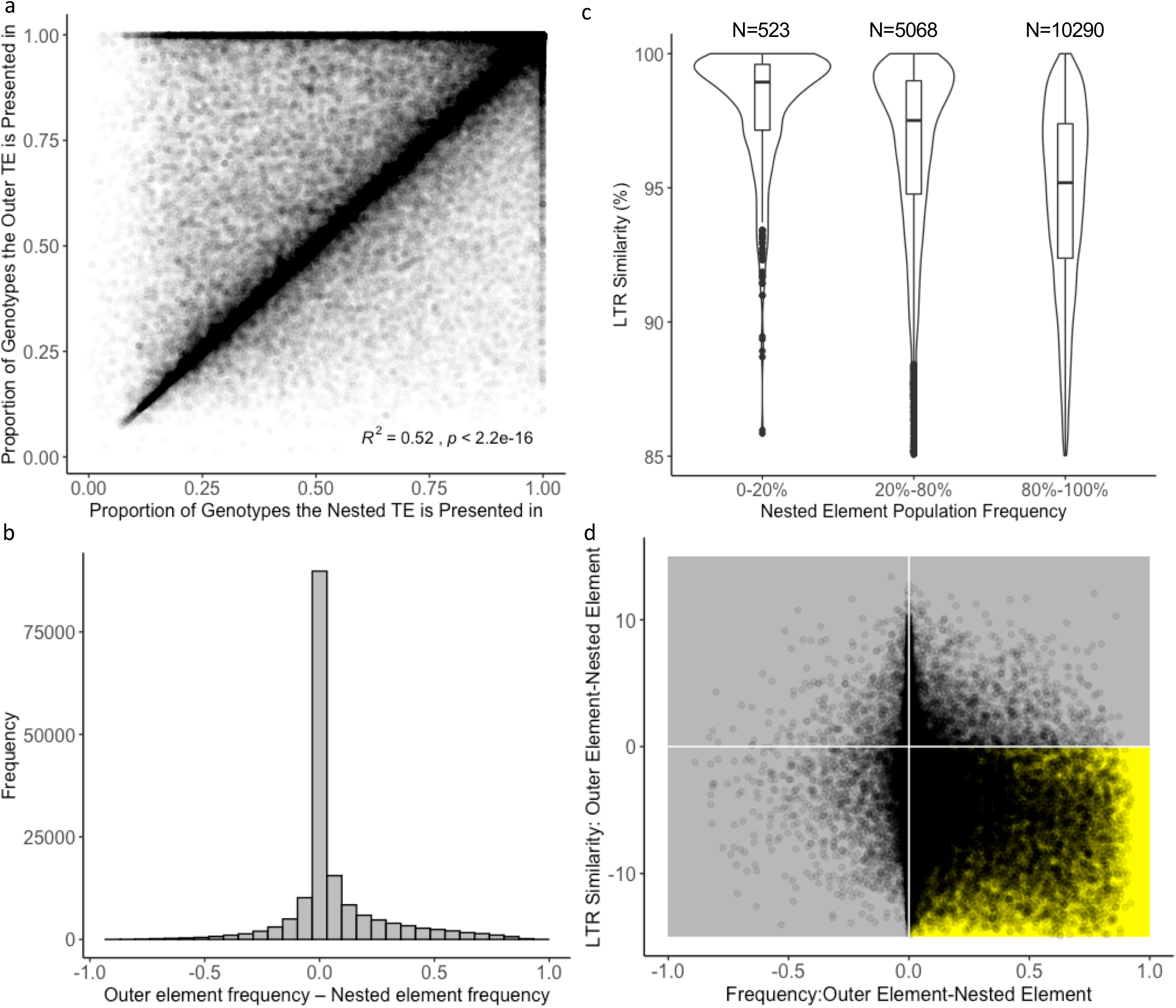
Relationship between population frequency of nested elements and the elements in which they are nested. a) Proportion of genotypes a TE is present in between nested TEs and the TE in which the element is nested. b) Distribution of the proportion of genotypes the outer TE is present in minus proportion of genotypes in which the nested TE is present. c) LTR similarity distributions for nested elements that are nested in TEs that are fixed or nearly fixed (frequency >0.95) in the population. This plot only contains nested LTRs as LTR similarity estimates are not available for other orders. d) Relationship between LTR similarity and frequency for outer elements minus nested elements. Points in the gold quadrant meet biological expectations that the outer element has a lower LTR similarity and is at higher frequency than the nested element. This plot only contains instances in which the outer and nested elements are both LTRs.

Within the set of outer elements that were fixed or nearly fixed in the population (frequency greater than 0.95), there was a continuous range of frequencies for inner elements that likely represent nested insertions with a range of ages. Using LTR similarity as a proxy to LTR age, we tested if there is a relationship between the LTR similarity of the nested element contained within fixed elements and their frequency in the population. This analysis could only be done for LTR nested elements as the other orders do not have an accurate metric to approximate age. As hypothesized, those nested elements that were at lower frequency in the population had higher LTR similarity than those that were at high frequency in the population (Figure 3c), and this was observed independent of the order of the outer element (Figure S7).

To further address this question of the timing of nested insertions into outer elements we focused on the subset of nested-outer element pairs in which both elements were LTRs and therefore had LTR similarity information. As with frequency, the quality of the LTR similarity metric was assessed with the expectation that the similarity of the nested insertions should be higher than that of the outer element. Across the LTR nesting pairs, 84.12% had a higher LTR similarity for the inner element, indicating an overall high quality of this data (Figure S8). Using the combination of frequency and LTR similarity we tested the extent to which an outer element and nested element insertion occurred at a similar time or weather the nested insertions occur over a range of time based on the LTR similarity (Figure 3d). The majority of nested-outer element pairs have nearly identical frequencies (Figure 3a,d), but with a range of LTR similarities. The most likely interpretation of these results is that in these cases the nested insertion occurred very near the timing of the outer element insertion and that the distribution of mutation accumulation is different for nested versus outer elements. This could be the result of methylation that occurs shortly after the initial insertion of the outer element. The paired outer and nested insertion will then insert as a unit in subsequent insertion events further perpetuating this relationship. While this finding is true for a large majority of the nested-outer pairs it should be noted that there are still a substantial number of pairs that have a range of frequency differences and LTR similarity differences that likely represent longer periods of time between the insertion of the outer element and the subsequent insertion of the nested element (Figure 3d).

### Relationship between location of TEs relative to genes and frequency in the population

TEs that are located in or near genes have the potential to change expression patterns of a gene (Hirsch and Springer 2017) or the product a gene encodes (Lisch 2013). In some cases, these functional insertions are a substantial distance from the gene, such as a 57kb upstream Harbinger-like DNA transposable element that represses *ZmCCT9* in cis and promotes flowering under long days (Huang *et al.* 2018). Others are much closer, such as a *STONER* element that inserted 42 bp upstream of the *Cg1* transcription start site and results in a chimeric fusion of the *STONER* element and the gene impacting the juvenile to adult vegetative transition (Chuck *et al.* 2007). Based on these prior studies, we sought to test genome-wide if the frequency and distribution of TE insertions varies based on their proximity and orientation with respect to genes in the genome. In order to test this, we first categorized TEs into categories using the following hierarchical ordering: gene completely within the TE, TE completely within the 5’ UTR, completely within the 3’ UTR, completely within an exon, completely within an intron, completely encompassed by a gene, 0-1 kb upstream of a gene, 1-5 kb upstream of a gene, 5-10 kb upstream of a gene, 0-1 kb downstream of a gene, 1-5 kb downstream of a gene, 5-10 kb downstream of a gene, and intergenic (greater than 10kb from the nearest gene).

There are relatively few gene proximal TEs that are actually within the gene body (orange in Figure 4a-c). Only 4.34% of LTR proximal TEs (1,640/37,838) and 3.79% of Helitron proximal TEs (334/9,056) are actually within the gene, while 11.52% (5,534/48,037) of TIR proximal TEs are within the gene body. In general those TEs that are within the gene body are at higher frequency than those that are outside of the gene body, and this is particularly true for those that not contained within an intron. For LTRs the age of the proximal TEs could also be evaluated, and those TEs that were within the gene body were also enriched for being younger based on LTR similarity than those proximal TEs that are outside of the gene body (Figure 4d; p-value < 0.001). This finding is somewhat unexpected given the potential deleterious functions that TE insertions could have on the expression of a gene and the integrity of the encoded protein, and likely indicates a positive effect of these insertions allowing them to rise in frequency relatively quickly.

**Figure 4.**
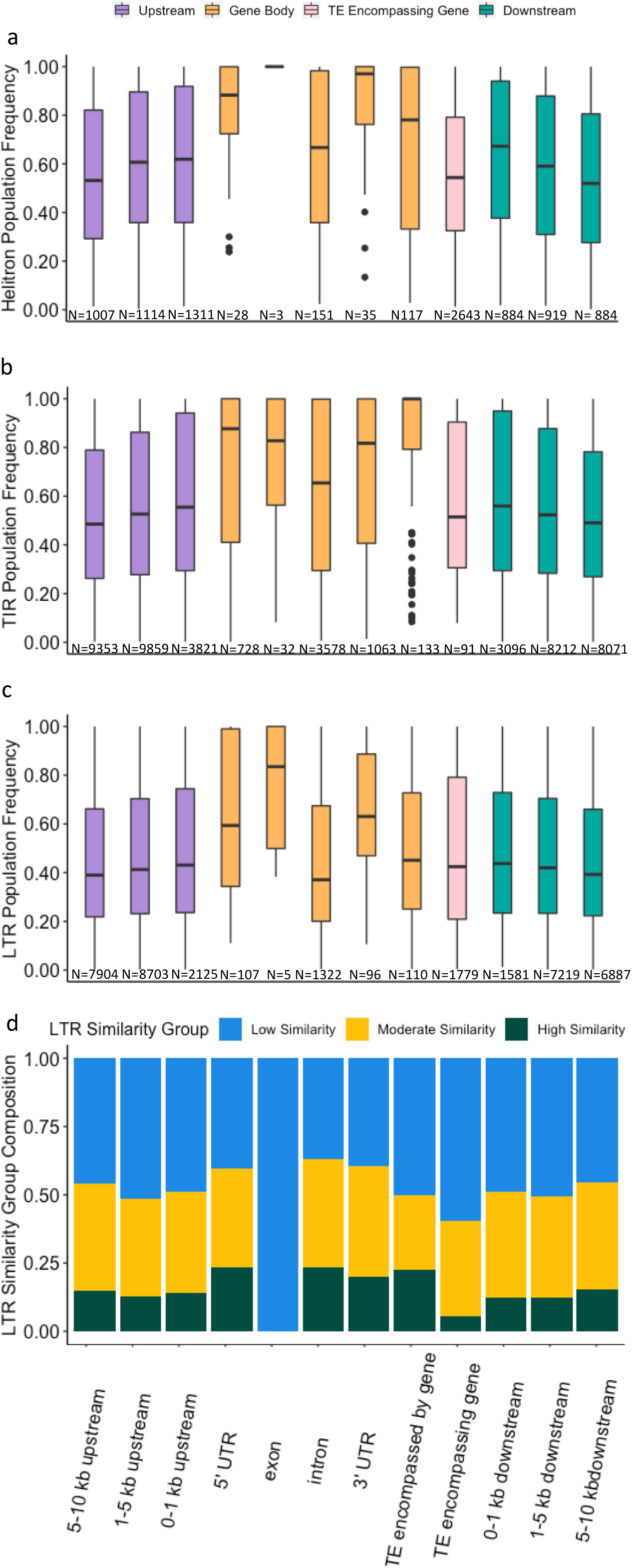
Relationship between TE frequency and relative position to the nearest gene. Helitrons (a), TIRs (b), and LTRs (c) were categorized using the following hierarchy: gene completely within the TE, TE completely within the 5’ UTR, completely within the 3’ UTR, completely within an exon, completely within an intron, completely encompassed by a gene, 0-1 kb upstream of a gene, 1-5 kb upstream of a gene, 5-10 kb upstream of a gene, 0-1 kb downstream of a gene, 1-5 kb downstream of a gene, 5-10 kb downstream of a gene, intergenic (not shown in figure). d) Proportion of LTRs with low LTR similarity (LTR similarity <95%), moderate LTR similarity (LTR similarity between 95-99%), and high LTR similarity (LTR similarity >99%) in each gene proximity category.

TEs that are proximal to a gene, but not contained within a gene, have much lower frequencies than was observed for TEs within gene bodies (purple and green in Figure 4a-c). The frequency further decays at increasing distance from the gene in both the 5’ and 3’ directions, and this is consistent across the three orders. Proximal TEs outside of the gene body also have a relatively higher proportion of insertions with low LTR similarity (older insertions) relative to those that are within genes (Figure 4d). The final class of proximal TEs, those that encompass a gene, show similar frequency to TEs that are proximal, but not within a gene, and are significantly depleted for insertions with high LTR similarity (p-value < 0.001; pink in Figure 4a-d). The genes contained within these TEs are all in the non-syntenic gene space relative to sorghum and rice. Genes in the non-syntenic gene space account for less phenotypic diversity than those in the syntenic gene space and are likely under different selective pressures in general compared to syntenic genes and TEs contained within syntenic genes (Schnable *et al.* 2011; Brohammer *et al.* 2018).

### Many TE insertional polymorphisms are not tagged by SNP markers

There is a limited number of plant species in which a species or population level cataloging of TE presence/absence variation has been conducted at a genome-wide level (Quadrana *et al.* 2016; Stuart *et al.* 2016; Carpentier *et al.* 2019; Chen *et al.* 2020; Dominguez *et al.* 2020). As such, the contribution of TEs to phenotypic variation, utility for genomic prediction, and other applications linking genotypes and phenotypes has only been minimally evaluated. In many instances, a linked marker has been identified as significant and with fine-mapping it is revealed that the causative variant is in fact a polymorphic TE insertion (e.g. (Studer *et al.* 2011)).

We sought to test the extent to which the extensive number of polymorphic TE insertions that were classified in this study were in linkage disequilibrium (LD) with genome-wide SNP markers to begin to understand the extent to which phenotypic variation that is caused by TEs has or has not been accounted for in previous studies that used only SNP markers. For this analysis, all SNPs within 1Mb plus or minus a TE were evaluated and the SNP that was in the highest LD was recorded. The majority of TEs were in moderate (r^2^ 0.5-0.9) to high (r^2^ >0.9) LD with a nearby SNP marker, but there was a subset of 49,382 (19.9%) TEs that were in low LD (r^2^ < 0.5) with all SNPs within a 1Mb window of the TE (Figure 5a). LTRs had the highest portion of TEs in high LD with a SNP (58.8%), while TIRs had 45.9% in high LD, and Helitrons had only 40.5% in high LD (Figure 5b). Of the subset of TEs that had a SNP in high LD (>0.9), the majority (86.50%) of the SNPs were within 200Kb of the TE (Figure 5c). This distance to the SNP with the highest LD increased when TEs in all levels of LD with SNPs were included (Figure S9). TEs that were at very high frequency within the population were generally in low LD with SNPs, and those that were at moderate frequency in the population were in moderate to high LD with SNPs (Figure 5d-f). This trend was consistently observed across orders of TEs.

**Figure 5.**
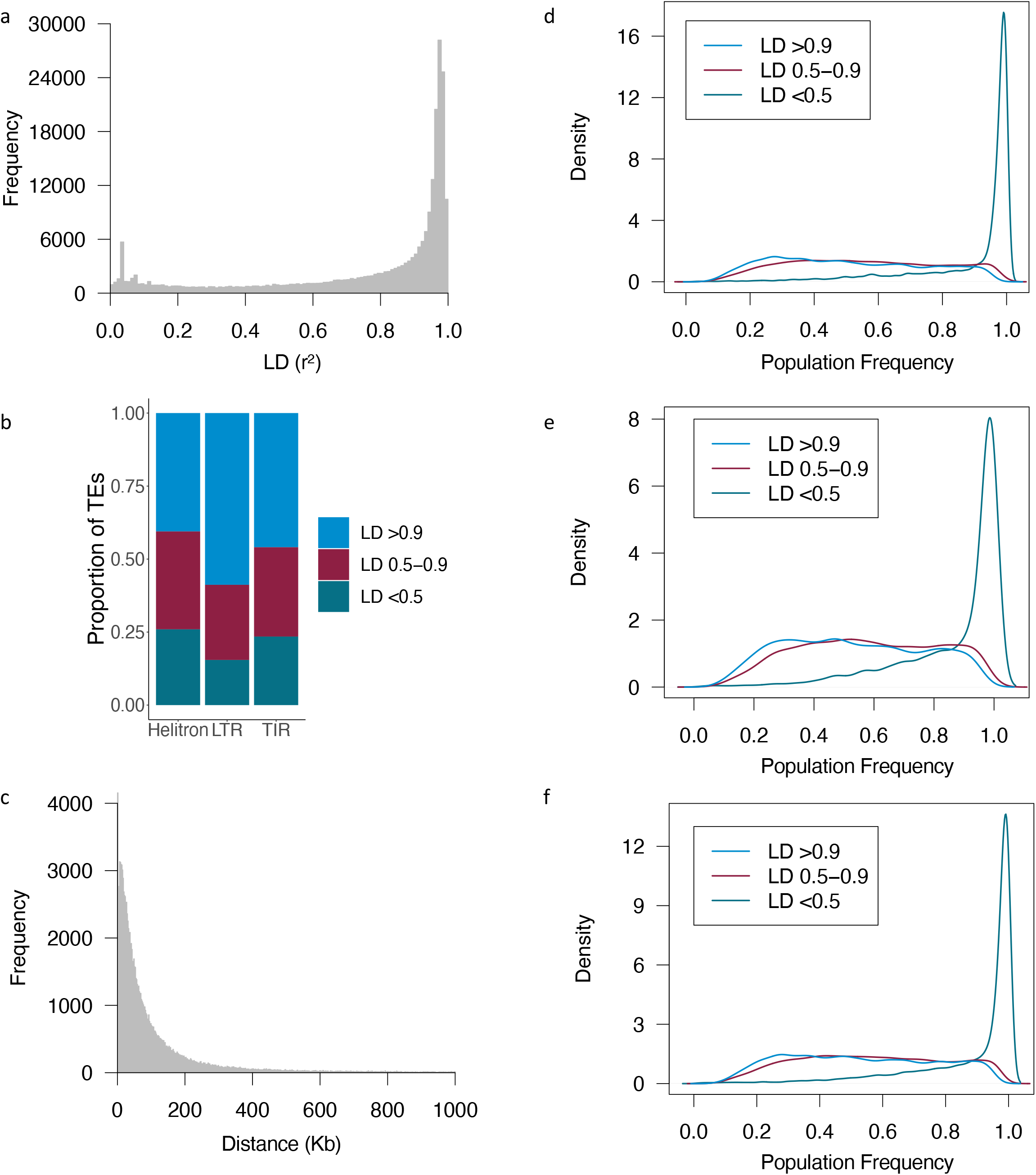
Linkage disequilibrium between TEs and SNPs in a panel of diverse inbred lines. a) Linkage disequilibrium (LD) between TEs and the SNP with the highest LD within 1Mb of the middle of the TE. b) Proportion of TEs in high (r^2^ > 0.9), moderate (r^2^ 0.5-0.9), and low (r^2^ <0.5) LD with SNPs within 1 Mb of the middle of the TE. Category is based on the SNP with the highest LD in the window. c) Distance between TEs and the SNP with the highest LD to it for TEs that had a SNP in high (r^2^ > 0.9) LD. Distance is calculated as the middle of the TE to the SNP. Only SNPs within 1 Mb of a TE were evaluated. d-f) Density plots of population frequencies for TEs in high, moderate, and low LD with SNPs based on the highest LD within 1Mb of the middle of the TE for LTRs (d), Helitrons (e), and TIRs (f). Only TEs with less than 25% ambiguous calls are included in these plots.

There are many metrics to assess LD and they represent different aspects of LD (Flint-Garcia *et al.* 2003; Slatkin 2008). For example, D’ measures only recombinational history, and r^2^ measures both recombinational and mutational history. We assessed LD between SNPs and TEs using D’ as the metric and observed substantially more TEs in near or perfect LD with a SNP (Figure S10a,b). Only 357 TEs had a D’ value of less than 0.9 to a SNP within 1 Mb of the TE, and a relationship with population frequency was no longer observed. The average distance between the SNP and TE decreased relative to r^2^ for the SNP in highest LD with a TE (102,838 for r^2^ vs. 50,371 for D’; Figure 5c and Figure S10c). This result makes sense as TE insertions represent different mutational events from the SNPs to which they are being evaluated and this is not reflected in the D’ metric.

Overall, while TE presence/absence patterns generally reflect maize breeding history (Figure S4), there are TE insertional polymorphisms that are not tagged by SNPs across different metrics (Figure 5 and Figure S10). These TEs that are not tagged by SNPs may be of important phenotypic consequence in maize, as has been shown in tomato (Dominguez *et al.* 2020) and rice (Akakpo *et al.* 2020). The high number of TEs in low to moderate LD with SNPs based on r^2^ is particularly important to this point, as r^2^ directly measures how different markers correlate with each other, and therefore how well a particular SNP would correlate with a potential causative TE. Including TE insertional polymorphism will likely be important in understanding the full complexity of phenotypic trait variation and local adaptation, and developing improved maize varieties in the future.

### Conclusion

TE insertional polymorphisms can play a crucial role in reshaping the phenotype of plants. In this study, we used the whole genome resequencing data, four high quality reference genomes with de novo annotated TEs, and random forest machine learning models, to generate a high-quality set of genome-wide insertional polymorphism data for 509 diverse maize lines. The majority of TE insertions (both nested and non-nested) were at the low to moderate frequency in the population. Within the LTR retrotransposon insertions, we observed a strong negative relationship between the frequency of the insertion in the population and the LTR similarity within the inserted element. Population frequency information coupled with LTR similarity also allowed us to determine that the majority of nested insertions (i.e. those insertions that are within another TE insertion) likely occur near the same time as the insertion of the outer element. Finally, analysis of LD between genome-wide SNP variants and TE insertional polymorphisms revealed that over 19.9% of TE insertional polymorphisms are not well tagged (R2 > 0.5 by nearby SNPs. This result has major implications when interpreting the results of genome-wide association studies (GWAS) that have been conducting using only SNP markers. Future work utilizing insertional polymorphism information may shed light into unexplained phenotypic variation in diverse germplasm such as this population.

## ACKNOWLEDGMENTS

This work was funded in part by NSF Grant IOS-1546727 to CNH and SEM, NSF Grant IOS-1934384 to CNH and NMS, and USDA Grant 2018-67013-27571 to CNH. The authors acknowledge the Minnesota Supercomputing Institute (MSI) at the University of Minnesota for providing resources that contributed to the research results reported within this paper. The authors would also like to thank Dr. Karin Dorman and Keting Chen (Department of Statistics and the Department of Genetics, Development and Cell Biology) at Iowa State University and Michael Burns (Department of Agronomy and Plant Genetics, University of Minnesota) for helpful discussion on the machine learning method used for TE present/absent calling.

## Notes

### Competing Interest Statement

The authors have declared no competing interest.

