## Supplementary material for "Whole Genome Variation of Transposable Element Insertions in a Maize Diversity Panel": Figures S1-10; Tables S1-4; Files S1-5: Figures_final.pdf

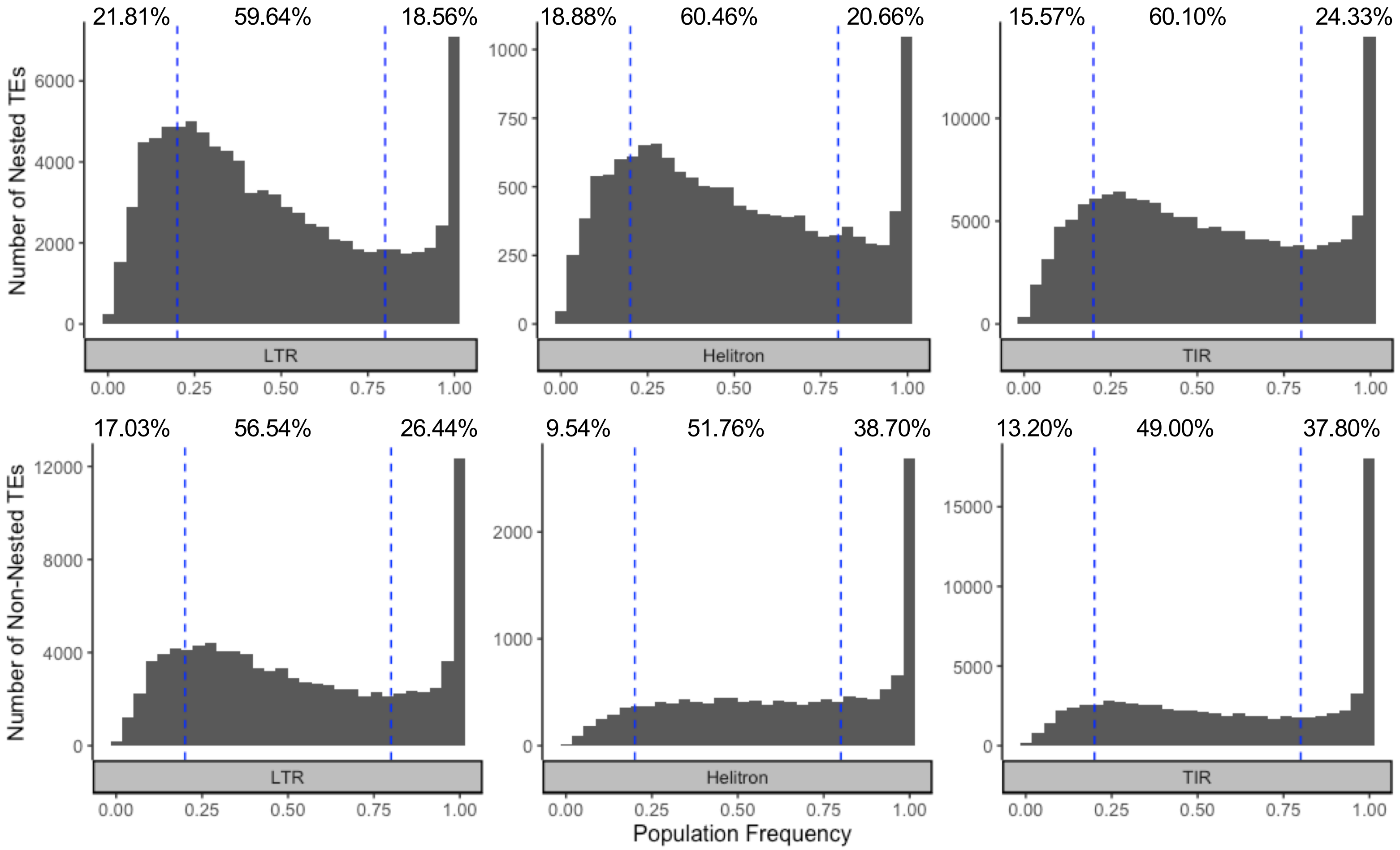


**Figure 1.** TE frequency distribution of a non-redundant set of TEs annotated in the B73, Mo17, PH207, and W22 genome assemblies. Short read sequence data from 509 genotypes were aligned to each genome assembly. Using a random forest machine learning method, TEs were classified into present (probability present >= 0.7), absent (probably present <=0.3), and all the other TEs were classified as ambiguous. For homologous TEs that were present in more than one assembly, the mean frequency across assemblies was calculated for the non-redundant set. TEs with less than 25% ambiguous calls are included (455,418 TEs). Percentages indicate the percent of low (<20%), moderate (20-80%), and high (>80%) frequency TEs in each order.

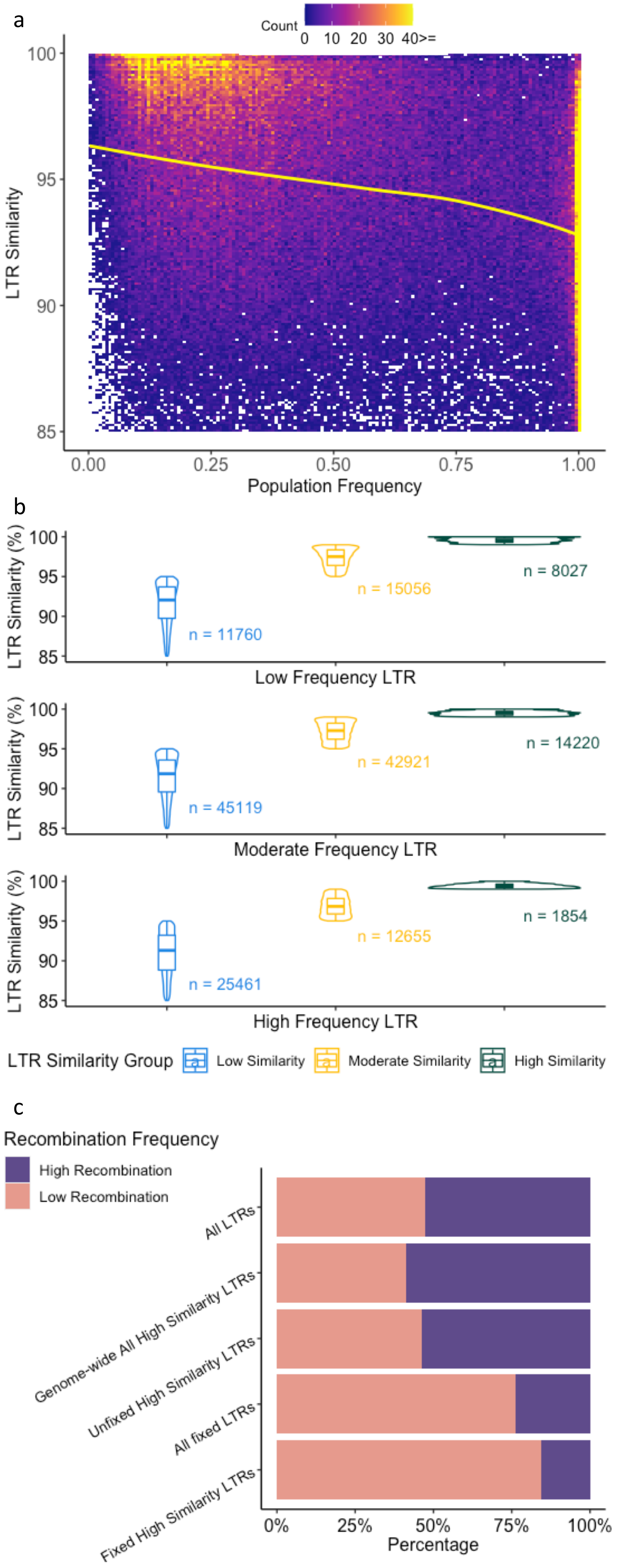


**Figure 2.** Relationship between TE similarity and frequency in a population of diverse inbred lines. a) Heatmap of LTR similarity versus frequency where white boxes indicate no TEs present at a particular frequency-by-LTR similarity. Yellow line is a LOESS curve fit through the data (n=177,073). b) Relationship of LTR similarity in categories of low similarity (LTR similarity <95%), moderate similarity (LTR similarity between 95-99%), and high similarity (LTR similarity >99%), and frequency in categories of low frequency (<20%), moderate frequency (20-80%), and high frequency (>80%). c) Proportion of different groups of LTRs in the low and high recombination portions of the genome based on B73 reference (n=108,968).

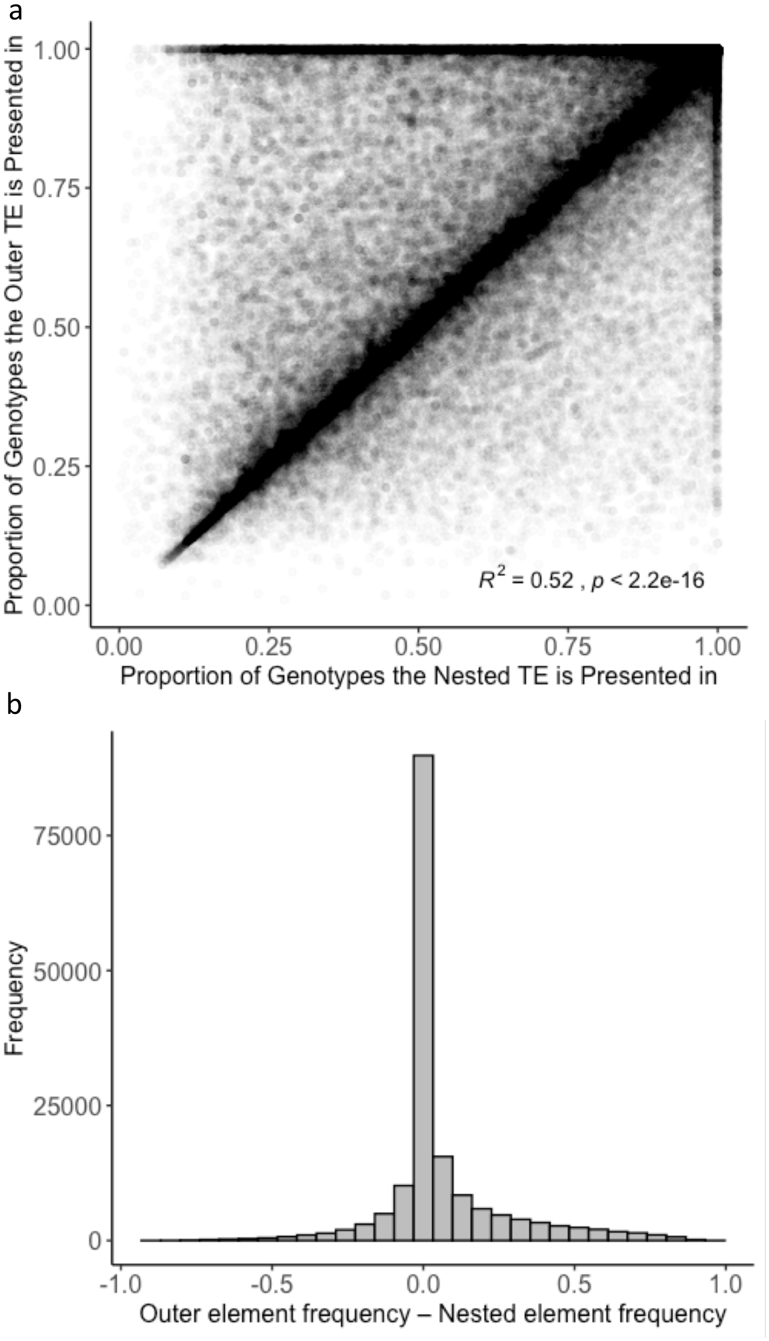


**Figure 3.** Relationship between population frequency of nested elements and the elements in which they are nested. a) Proportion of genotypes a TE is present in between nested TEs and the TE in which the element is nested. b) Distribution of the proportion of genotypes the outer TE is present in minus proportion of genotypes in which the nested TE is present. c) LTR similarity distributions for nested elements that are nested in TEs that are fixed or nearly fixed (frequency >0.95) in the population. This plot only contains nested LTRs as LTR similarity estimates are not available for other orders. d) Relationship between LTR similarity and frequency for outer elements minus nested elements. Points in the gold quadrant meet biological expectations that the outer element has a lower LTR similarity and is at higher frequency than the nested element. This plot only contains instances in which the outer and nested elements are both LTRs.

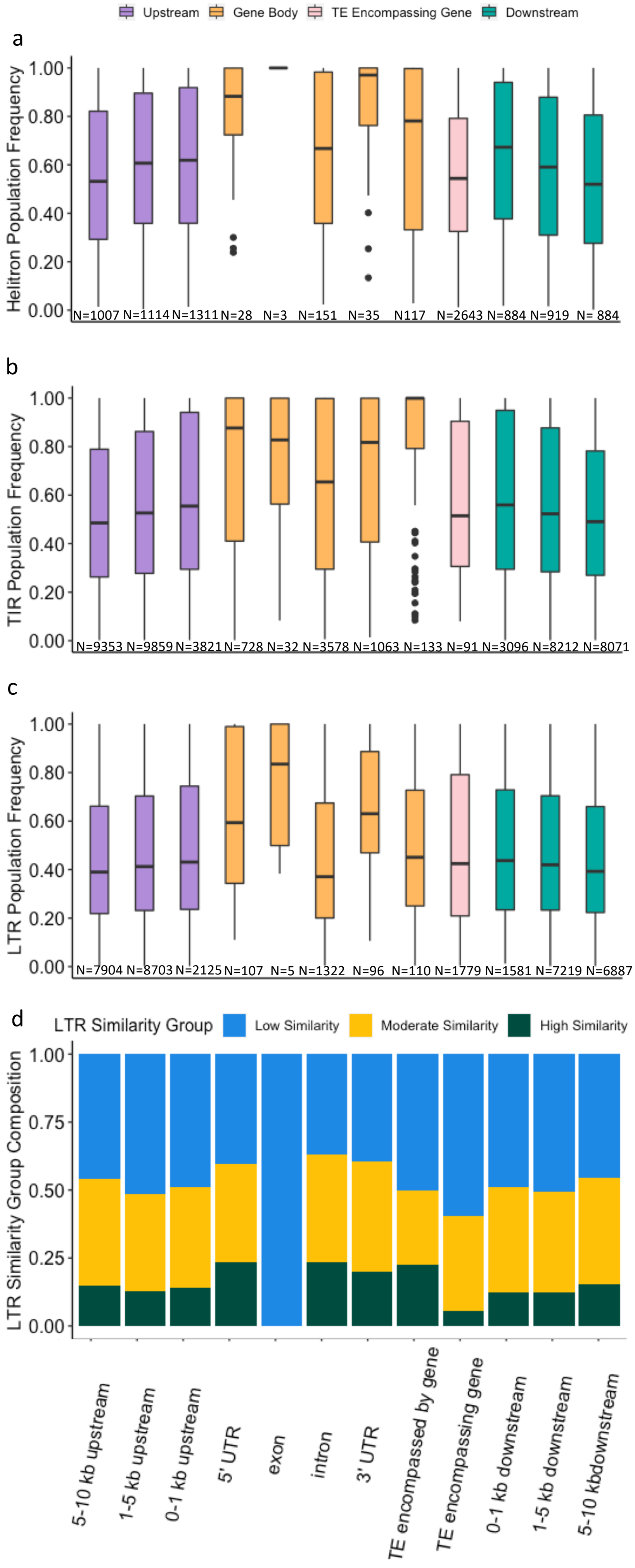


**Figure 4.** Relationship between TE frequency and relative position to the nearest gene. Helitrons (a), TIRs (b), and LTRs (c) were categorized using the following hierarchy: gene completely within the TE, TE completely within the 5' UTR, completely within the 3' UTR, completely within an exon, completely within an intron, completely encompassed by a gene, 0-1 kb upstream of a gene, 1-5 kb upstream of a gene, 5-10 kb upstream of a gene, 0-1 kb downstream of a gene, 1-5 kb downstream of a gene, 5-10 kb downstream of a gene, intergenic (not shown in figure). d) Proportion of LTRs with low LTR similarity (LTR similarity <95%), moderate LTR similarity (LTR similarity between 95-99%), and high LTR similarity (LTR similarity >99%) in each gene proximity category.

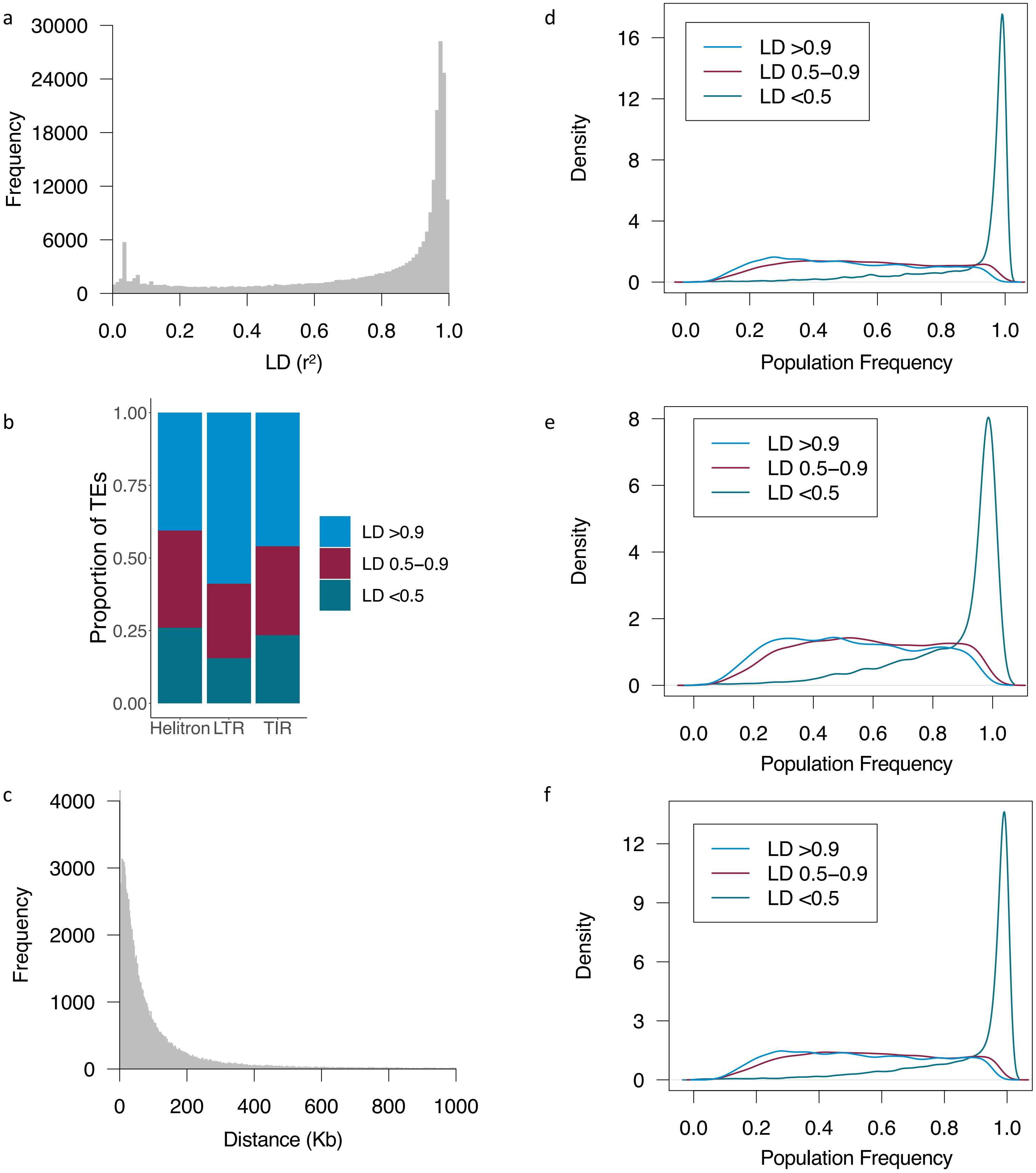


**Figure 5.** Linkage disequilibrium between TEs and SNPs in a panel of diverse inbred lines. a) Linkage disequilibrium (LD) between TEs and the SNP with the highest LD within 1Mb of the middle of the TE. b) Proportion of TEs in high ($r^2$ > 0.9), moderate ($r^2$ 0.5-0.9), and low ($r^2$ <0.5) LD with SNPs within 1 Mb of the middle of the TE. Category is based on the SNP with the highest LD in the window. c) Distance between TEs and the SNP with the highest LD to it for TEs that had a SNP in high ($r^2$ > 0.9) LD. Distance is calculated as the middle of the TE to the SNP. Only SNPs within 1 Mb of a TE were evaluated. d-f) Density plots of population frequencies for TEs in high, moderate, and low LD with SNPs based on the highest LD within 1Mb of the middle of the TE for LTRs (d), Helitrons (e), and TIRs (f). Only TEs with less than 25% ambiguous calls are included in these plots.
