## Supplementary material for "Whole Genome Variation of Transposable Element Insertions in a Maize Diversity Panel": Figures S1-10; Tables S1-4; Files S1-5: supplemental_figures_final.pdf

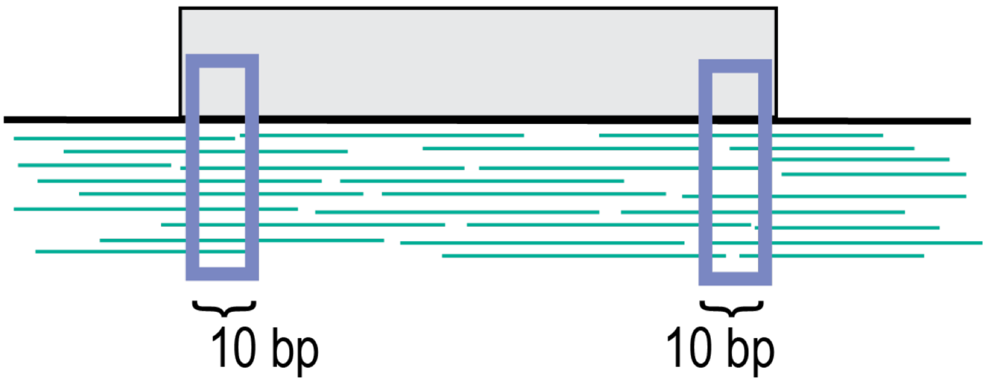


**Figure S1.** Schematic of windows used to get start and end coverage.

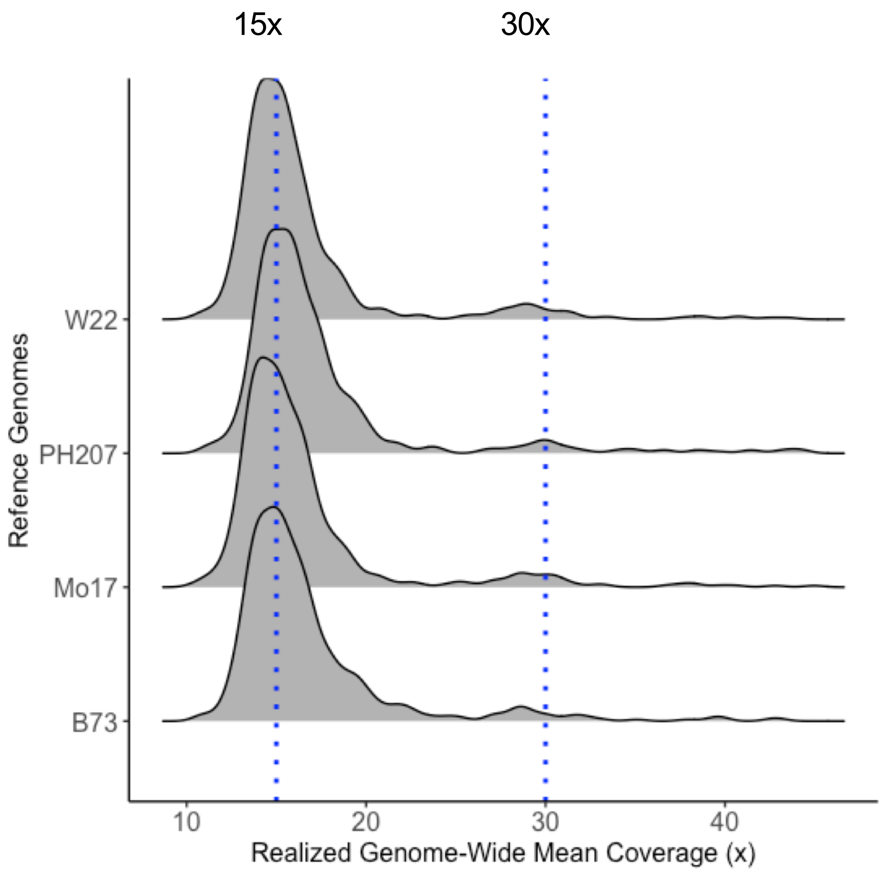


**Figure S2.** Realized genome-wide mean coverage for samples used in this study.

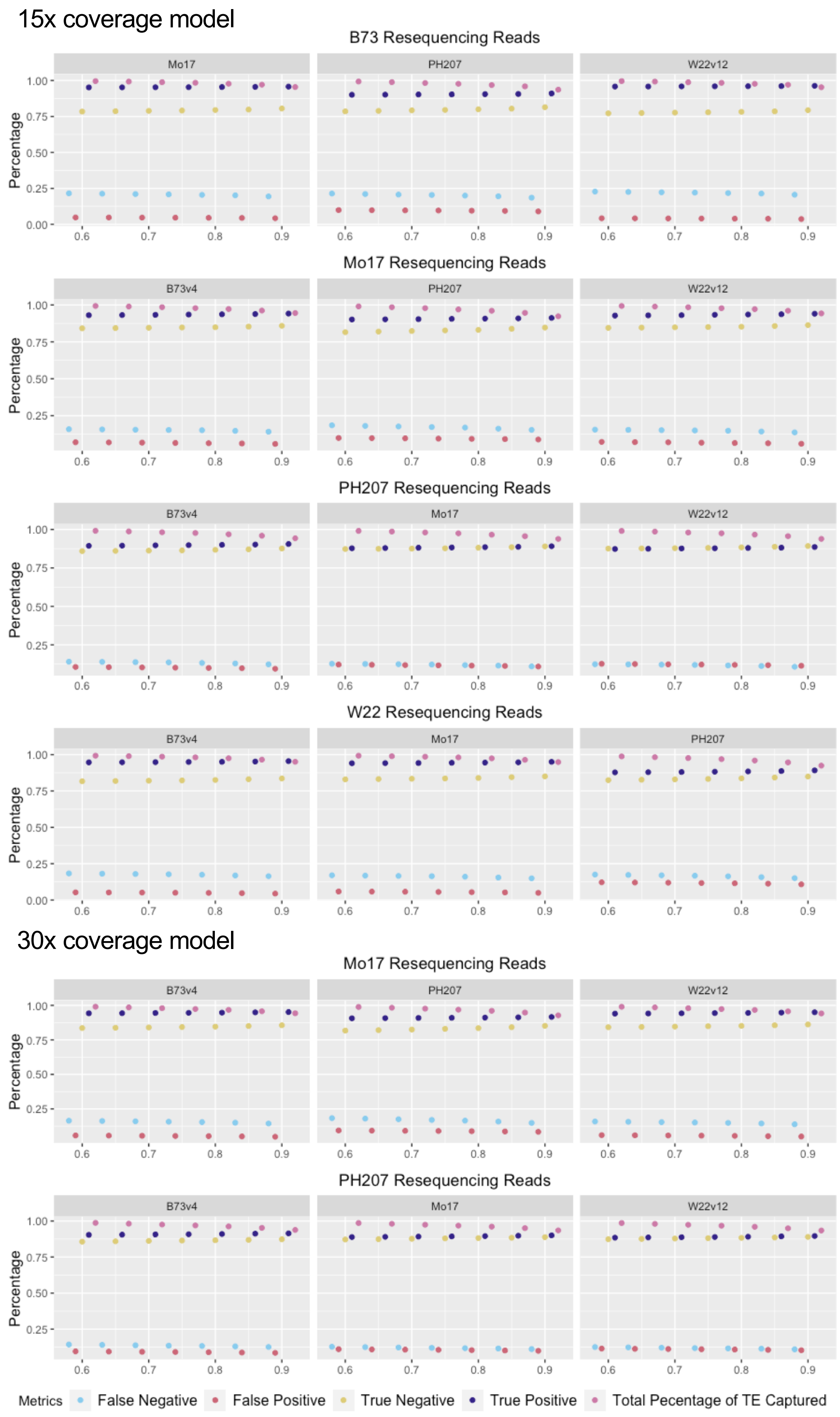


**Figure S3.** TE calling accuracy for the random forest machine learning model. Two models (one trained on features from 15x coverage samples and one trained on features from 30x coverage samples) were built and tested for false negative, false positive, true positive, true negative based on comparison of a gold standard set of TE polymorphisms from whole genome comparisons, and total percentage of TE captured for each test set.

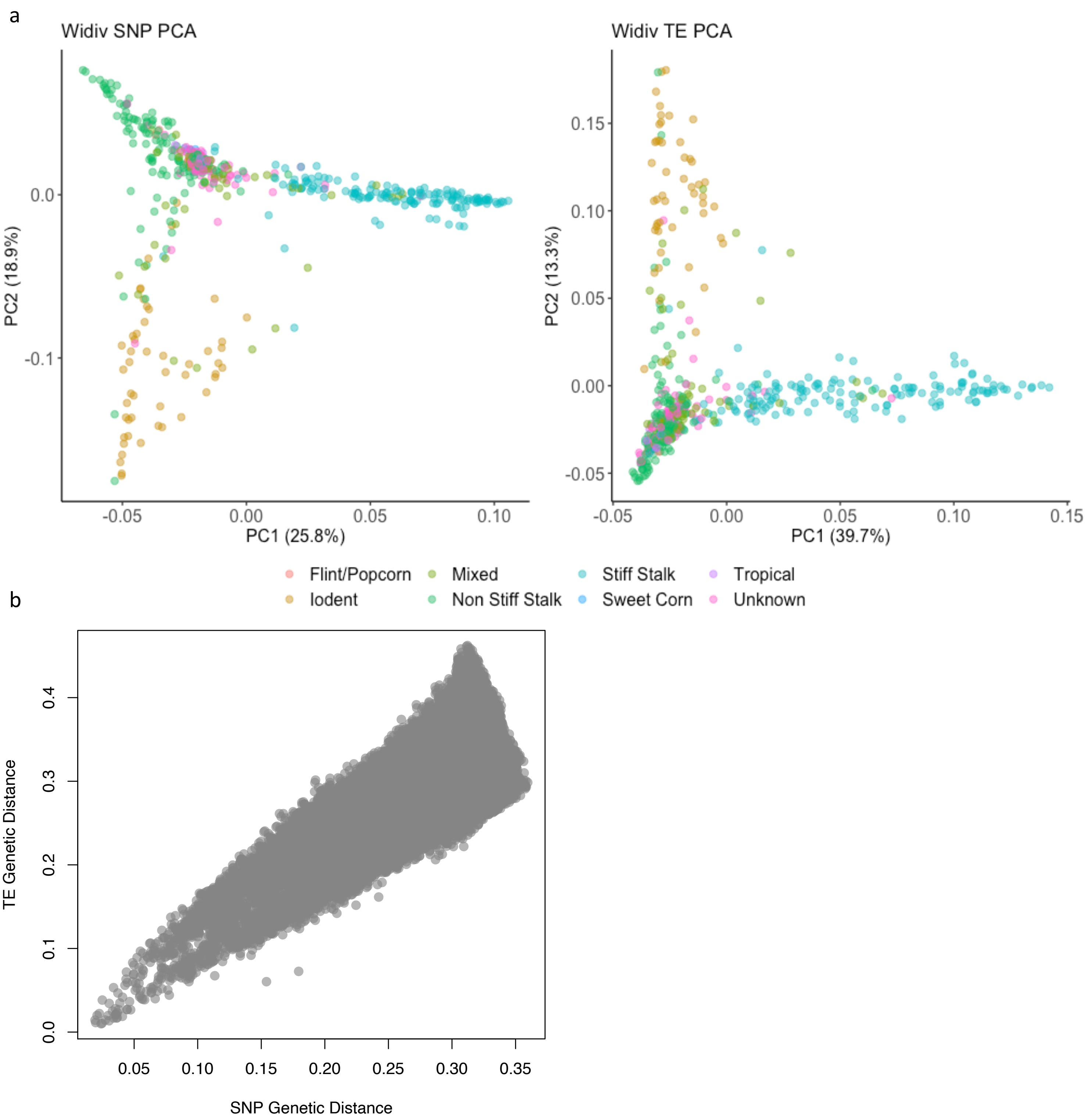


**Figure S4.** Genetic relationships based on different sources of variation in the genome. a) Principle component analysis (PCA) of SNPs (left) and TE presence/absence (right) in a set of diverse maize inbred lines. b) Relationship between pair-wise genetics distances determine using genome-wide SNP markers versus genome-wide TE presence/absence classifications. TEs and SNPs were called relative to the B73 reference genome assembly.

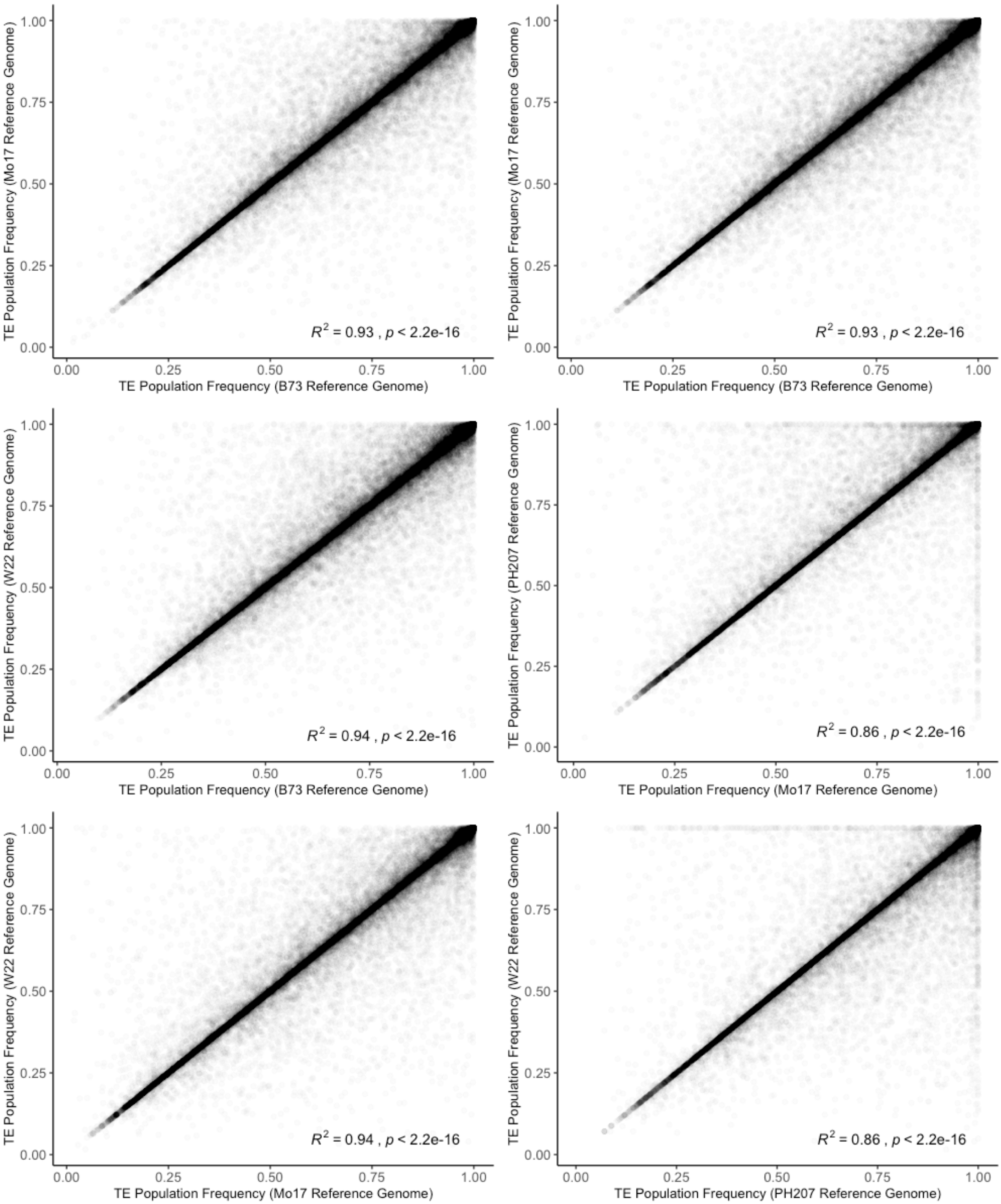


**Figure S5.** Proportion of genotypes a TE was called as 'Present' in between homologous TEs in each pair-wise comparison of genomes.

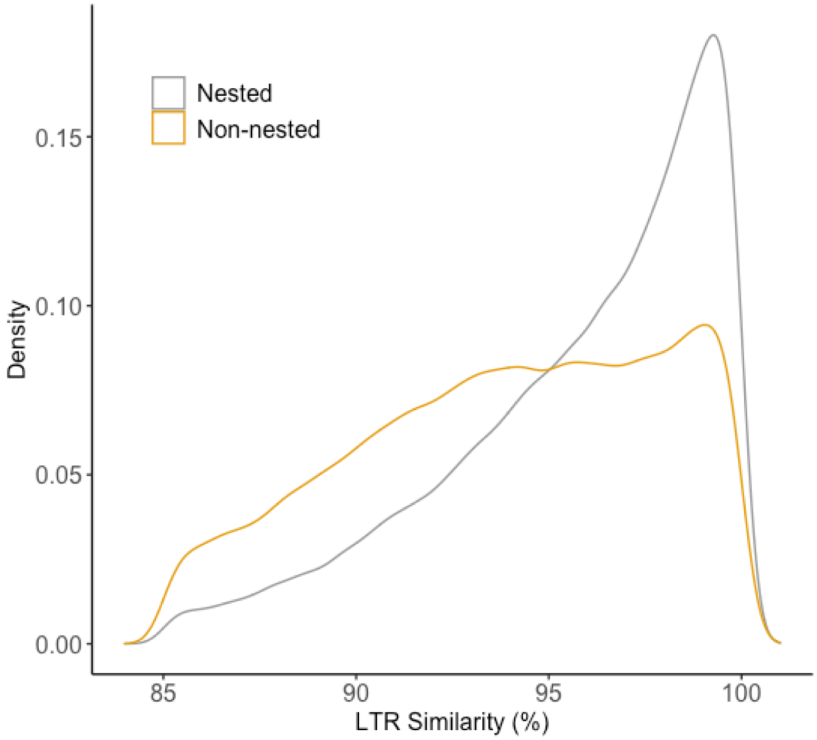


**Figure S6.** LTR similarity distribution of nested and non-nested elements in a non-redundant set of 177,073 LTR elements.

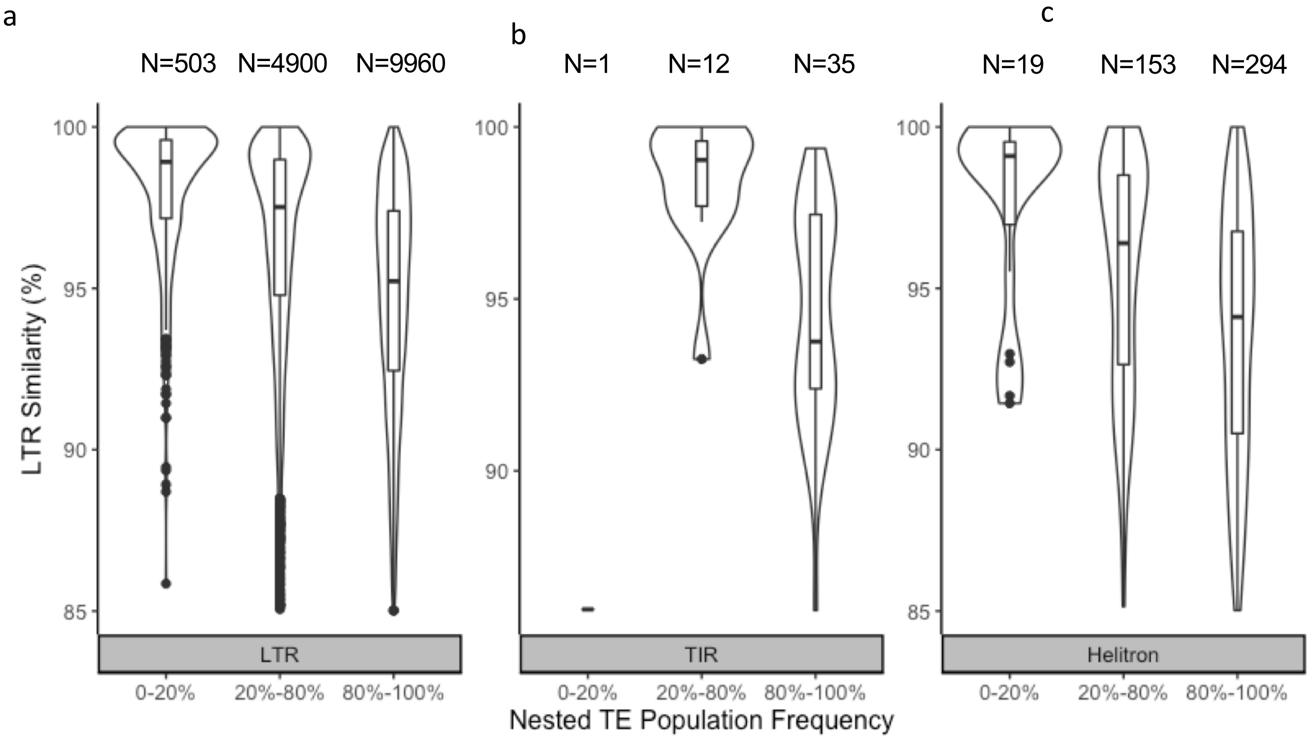


**Figure S7.** LTR similarity distributions for nested elements that are nested in TEs that are fixed or nearly fixed (frequency >0.95) in the population. Frequency distribution are broken down by the order of the outer element with a) LTR outer element, b) TIR outer element, and c) Helitron outer element. This plot only contains nested LTRs as TE similarity estimates are not available for other orders.

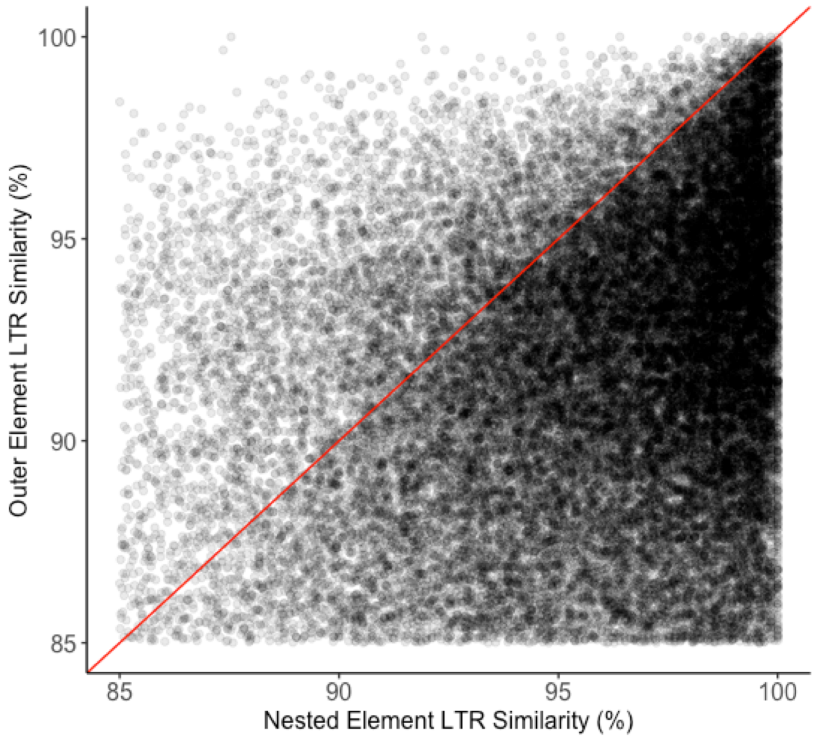


**Figure S8.** Relationship between LTR similarity of nested elements and the elements in which they have inserted. This plot only contains LTRs for the nested and outer elements as similarity estimates are not available for other orders. Data points that fall below the red line are instances where the nested element has a higher similarity than the outer element.

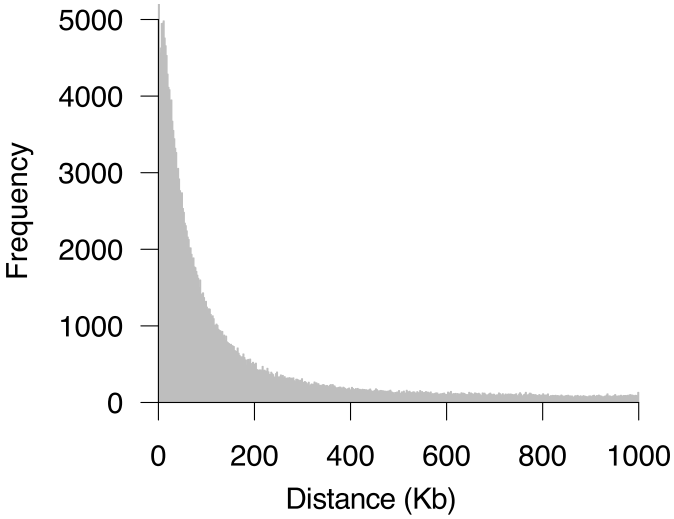


**Figure S9.** Distance between TEs and the SNP with the highest LD to it. Distance is calculated as the middle of the TE to the SNP. Only SNPs within 1Mb of a TE were evaluated.

a

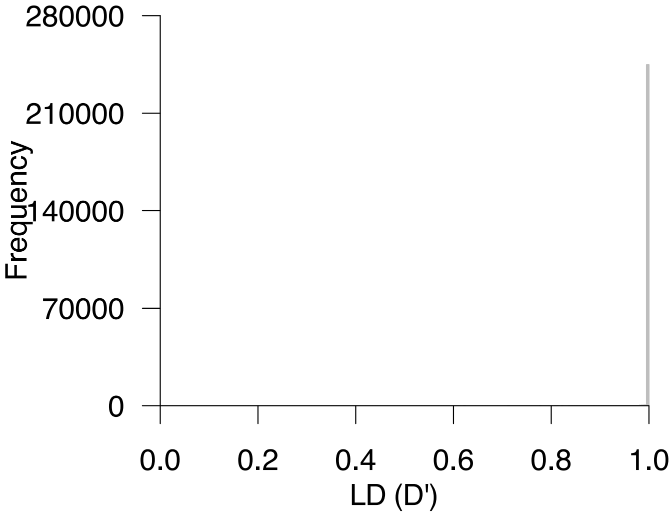


b

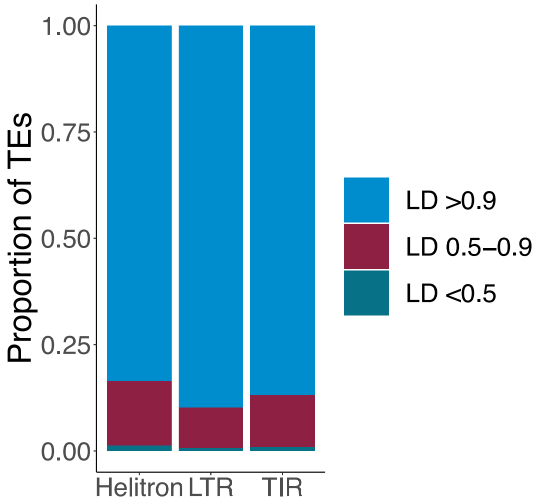


c

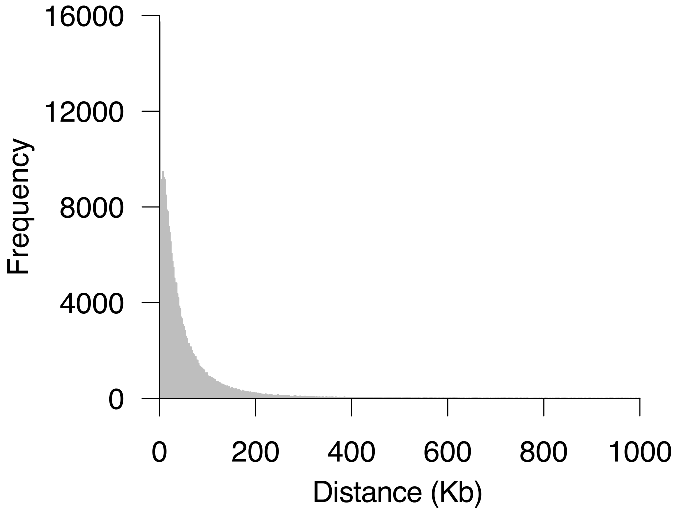


**Figure S10.** Linkage disequilibrium between TEs and SNPs in a panel of diverse inbred lines using D'. a) Linkage disequilibrium (LD) between TEs and the SNP with the highest LD within 1Mb of the middle of the TE. b) Proportion of TEs in high (D' > 0.9), moderate (D' 0.5-0.9), and low (D' <0.5) LD with SNPs within 1 Mb of the middle of the TE. Category is based on the SNP with the highest LD in the window. c) Distance between TEs and the SNP with the highest LD to it for TEs that had a SNP in high (D' > 0.9) LD. Distance is calculated as the middle of the TE to the SNP. Only SNPs within 1 Mb of a TE were evaluated. d) Density plot of population frequencies for TEs in high, moderate, and low LD with SNPs based on the highest LD within 1Mb of the middle of the TE.
